# Horizon: CNV interpretation through rapid automated ACMG-aligned pathogenicity analysis

**DOI:** 10.1101/2024.05.27.596089

**Authors:** Mariam Eldesouky, Suhana Shiyas, Amirul Islam, Md. Nahid Hasan, Md. Tanvir Hossain, Hosneara Akter, Bakhrom K. Berdiev, Wolfgang M. Kuebler, Proton Rahman, Marc Woodbury-Smith, Syed M. Hasan, Nasna Nassir, Mohammed Uddin

## Abstract

**Purpose:** Our study assesses the Horizon model, a novel CNV classification tool developed in line with American College of Medical Genetics (ACMG) guidelines, to enhance the classification of pathogenicity in CNVs.

**Methods:** Horizon utilizes a ranking-based algorithm, incorporating multiple proprietary databases and variant inheritance models as per ACMG standards. The model’s effectiveness was verified through Area Under the Curve (AUC) analyses on three datasets comprising 696 pathogenic inherited or *de novo* variants, as classified by clinical geneticists and several established tools.

**Results:** Horizon achieved an AUC of 0.97 in the discovery cohort, demonstrating high accuracy in CNV interpretation and proficiency in predicting pathogenicity. We observed an AUC of 0.87 in the *de novo* variant cohort and an overall AUC of 0.94 across all cohorts, surpassing tools like ClassifyCNV and AnnotSV. It showed particular effectiveness in interpreting duplication CNVs and the highest performance for CNVs sized 3-5 Mb.

**Conclusion:** The Horizon model offers robust and accurate CNV interpretation, outperforming existing tools and aligning closely with clinical evaluations. Its comprehensive approach, integrating a range of genomic features and following ACMG guidelines, makes it a crucial tool in the genomic interpretation landscape, facilitating the rapid and accurate diagnosis of genetic disorders.

## Introduction

Copy number variations (CNVs) are a class of genetic variations, broadly categorized under structural variations (SVs), which play a key role in genomic diversity and disease etiology. CNVs are typically characterized by deletions or duplications of genomic segments of at least 50 bp in length. CNVs constitute a significant portion of the human genome, estimated to range between 4.8% and 9.5%.^1^ The effects of CNVs span a wide spectrum, ranging from benign phenotypic outcomes to considerable contributions in the onset of chronic debilitating diseases. These variations, especially when they alter gene coding segments, can significantly affect gene dosage and expression, leading to various syndromes.^2,3^ Their association with Mendelian disorders such as Potocki-Lupski syndrome and Williams-Beuren syndrome, neurodevelopmental disorders such as autism spectrum disorder (ASD) and schizophrenia, and various cancers,^4^ highlights the necessity to understand CNVs for comprehensive genomic analysis. Although technologies are evolving very rapidly to detect the full spectrum of structural variations (SVs), the annotation and classification of CNVs to determine their pathogenic potential remain critical for their practical application in clinical settings.

CNVs are classified into pathogenic, likely pathogenic, variant of unknown significance (VUS), likely benign, or benign, as per guidelines of the American College of Medical Genetics and Genomics (ACMG). These consider various aspects of a variant, including genomic content, predicted functional impact, and evidence in literature. Clinical Genome Resource (ClinGen) and DECIPHER pose a significant step towards standardizing CNV interpretation.^5–7^ However, characterizing most CNVs, which are unique and not recurrent, remains challenging due to the reliance on individual proficiency and extensive knowledge of clinical genetics.

To address these challenges, we introduce Horizon, an automated CNV classification tool aligning with the latest ACMG guidelines. Horizon leverages extensively populated databases for pathogenicity analysis and has demonstrated performance comparable to manual assessments by clinical evaluators. Furthermore, we compared Horizon’s sensitivity in detecting clinically relevant variants against two other CNV annotation tools following ACMG classification: ClassifyCNV and AnnotSV.^8,9^ Our findings suggest that Horizon is a highly promising tool that could substantially aid both research and clinical diagnostics in the realm of genetic variations.

## Materials and Methods

### Study design and data collection

To assess the performance of the Horizon model under various conditions, we compiled a total of 696 reported pathogenic CNVs and defined three distinct cohorts for further assessment. All CNVs were unified and remapped to the GRCh38/hg38 reference genome using the UCSC Genome Browser LiftOver tool.^10^ We did not include any studies that are part of Horizon’s internal databases. Since VUS interpretation can be extremely subjective and vary between studies, our analysis quantifies the precision of Horizon’s ability to recall these known pathogenic variants. For our accuracy, sensitivity, and recall rate, we have combined both “Likely Pathogenic” and “Pathogenic” into one single category called “Pathogenic”.

#### Discovery Cohort

This cohort comprised 212 patients with neurodevelopmental disorders, including epilepsy, ASD, hypotonia, speech and language disabilities. These patients were referred for whole genome microarray analysis after consultations with multiple neurologists. Chromosomal microarray analysis (CMA) was performed using the Illumina Global Screening Array-24 + v1.0, applying Illumina SNP genotyping technology and the Illumina CNVPartition 3.2.1 plug-in of GenomeStudio. Clinical geneticists reported pathogenicity of 43 variants (all of them validated by droplet digital PCR with TaqMan Assay) from a set of 1,766 identified CNV’s.^11^ We have termed this the discovery cohort due to the rigorous assessment of the variants by a single clinical geneticist and the use of one microarray platform. This cohort consists of relevant clinical primary pathogenic variants.

#### Literature Cohort

The literature cohort is a curation of over 491 pathogenic CNVs associated with various rare diseases (Table S1). These variants were collected through a systematic review of open-source publications using keywords such as “CNV”, “SV”, “Pathogenic”, “Clinical genome sequencing”, and “Whole Genome Microarray”. A significant portion of these variants were classified as “Pathogenic” or “Likely Pathogenic” by numerous clinical geneticists across the globe. A major caveat of this dataset is the use of different microarray and sequencing technologies, including old BAC arrays comprising highly inaccurate CNV breakpoints, leading to inaccurate pathogenicity classification.^12^ We have also obtained a dataset of 94 CNVs evaluated by two ACMG certified clinical geneticists.^8^ Given the discrepancies in variant interpretation between the clinical geneticists, we treated each clinician’s evaluation as a separate dataset and evaluated Horizon’s predictions for comparison.

#### *de novo* Cohort

We collated a secondary dataset of 162 pathogenic CNVs obtained from multiple studies, incorporating heterogeneous variants identified across various rare disorders and ethnicities. Sources included published data on congenital heart disease,^13^ cerebral palsy,^14^ and neurodevelopmental disorders.^15,16^

### Horizon model

Horizon employs a scoring system based on ACMG guidelines, classifying variants into five categories: Pathogenic, Likely Pathogenic, VUS, Likely Benign, and Benign. Horizon implemented multiple analytical modalities within their prediction models, including proprietary databases and public databases (i.e. Clinvar, DECIPHER, 1000k genomes). Horizon’s ranking model is based on the ACMG guideline defined for rare diseases.^5,17^

Horizon deploys multiple proprietary databases, and the algorithm utilizes these to assess variant inheritance, effect and function. A population control genome wide SV database comprising a large number of samples with different ethnicities allows the algorithm to filter common variants. The Horizon disorder database consists of a large number of curated clinically relevant (pathogenic, likely pathogenic) structural variants from peer reviewed published research papers on rare diseases and germline pan cancer. These databases were curated by the work of expert geneticists. The Horizon algorithm integrates a set of validated and ACMG supported genes known to cause different autosomal recessive or dominant rare diseases due to SVs. The curation of these proprietary databases is a multi-year effort by GenomeArc Inc. The score ranking of each variant for a patient is assessed based on the multivariate analysis that fulfills the pathogenicity requirements stated by the ACMG guidelines. The score was then converted into one of the five textual categories defined earlier.

### Comparison with other tools

To benchmark Horizon, we compared its performance with ClassifyCNV and AnnotSV.^8,9^ These tools, also based on ACMG guidelines, provide variant classifications used to plot ROC graphs and measure accuracy through AUC scores. All cohorts were annotated by ClassifyCNV, AnnotSV, and Horizon, and their interpretations were compared against initial classifications. In the present comparison we only considered “Pathogenic” variants; VUS were excluded due to the variation in interpretation that is unfortunately characteristic of such variants. For all our tests of the Horizon model, we applied four population control frequencies (0.01, 0.05, 0.001, and 0.005) to assess the improvement of AUC.

## Results

In our initial evaluation using the discovery cohort of 43 pathogenic CNVs, Horizon identified 41 as pathogenic and 2 as variants of uncertain significance (VUS), demonstrating the model’s precision. Receiver operating characteristic (ROC) curve analysis identified 0.005 as the optimal population control frequency for pathogenicity prediction. Subsequent use of this frequency to additional cohorts further validated its efficacy. Horizon’s ROC analysis in the discovery cohort yielded a high area under the curve (AUC) score of 0.97, surpassing AnnotSV and ClassifyCNV with AUCs of 0.86 and 0.78, respectively (Figure 1A). Importantly, Horizon maintained its conservative stance by not reclassifying any pathogenic CNVs as benign, unlike AnnotSV, which reclassified one. Sensitivity analysis comparing deletion and duplication CNVs revealed robust performance with AUCs of 0.95 and 0.99, respectively (Figure 1B). Size-based analysis across CNVs highlighted consistent accuracy (AUC > 0.9) in all categories, except for those measuring 1-3 Mb, which showed a slightly lower accuracy (AUC = 0.88) (Figure 1C).

**Figure 1.**
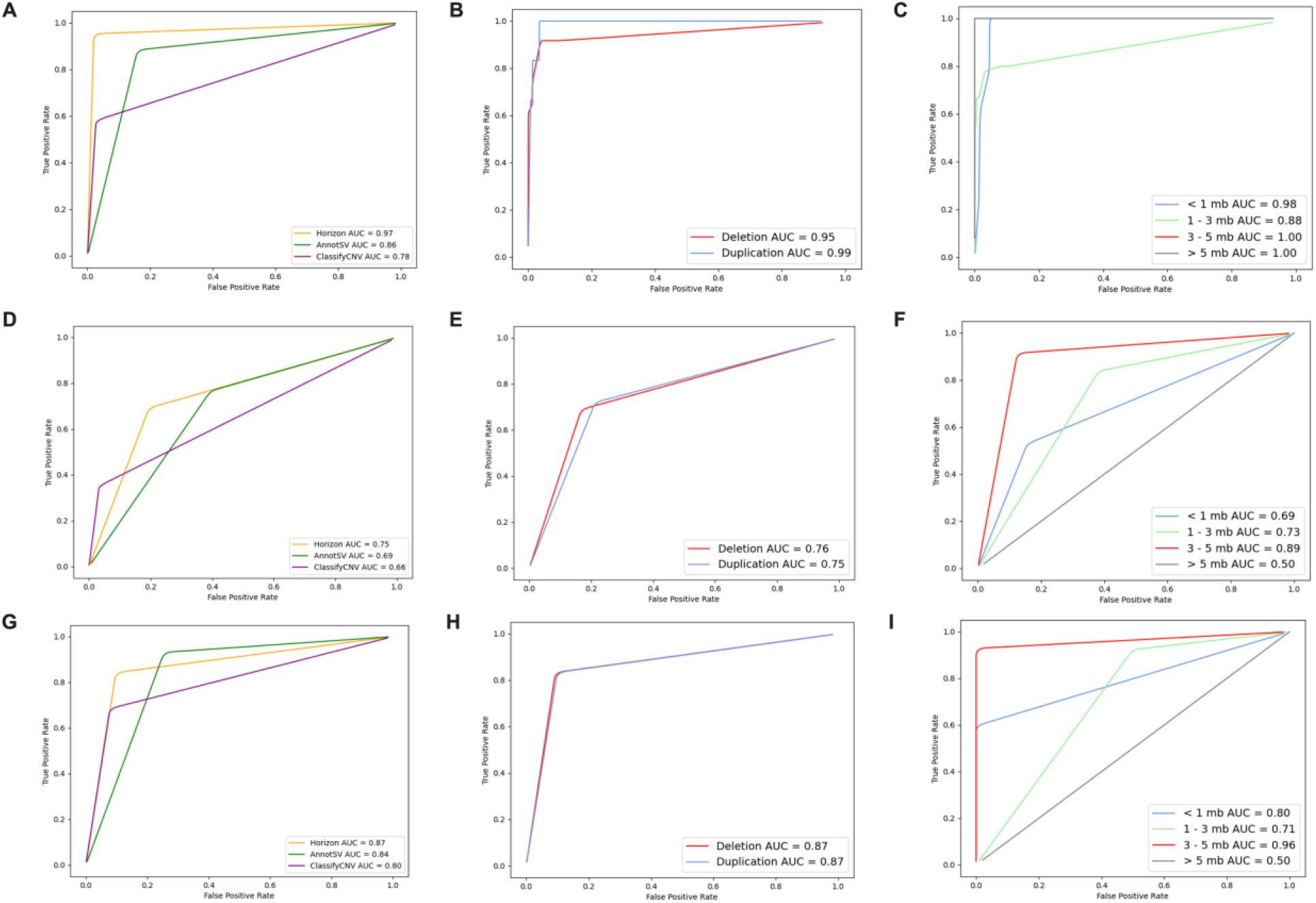
ROC analyses of the Discovery Cohort. **A**. Performance comparison of Horizon (yellow line) with AnnotSV (green line) and ClassifyCNV (purple line) for the discovery cohort; **B**. The CNV types, deletion (red line), duplication (blue line) and **C**. The different length of CNVs color coded ROC curves and their corresponding AUC. **D-F**. Literature cohort and Horizon CNV pathogenicity prediction AUC estimation compared to different CNV pathogenicity prediction tools, CNV types and different CNV lengths, respectively. **G-I**. *de novo* cohort and Horizon CNV pathogenicity prediction AUC estimation compared to different CNV pathogenicity prediction tools, CNV types and different CNV lengths, respectively.

For the literature cohort of 491 pathogenic variants, Horizon classified 343 as pathogenic and 148 as VUS, achieving an overall AUC of 0.75. This performance was superior to both AnnotSV (AUC = 0.69) and ClassifyCNV (AUC = 0.66) (Figure 1D). Horizon maintained strict classification standards, not reclassifying any of the 491 pathogenic variants as benign. In contrast, AnnotSV and ClassifyCNV identified 9 and 5 pathogenic variants as benign, respectively. In this cohort, the highest accuracy was observed among CNVs sized 3-5 Mb (AUC = 0.89), although accuracy decreased substantially for variants over 5 Mb (AUC = 0.5) (Figure 1F). Horizon’s prediction on the dataset interpreted by two ACMG evaluators showed some discrepancies, yielding an AUC of 0.70 and 0.74 with closer alignment with the second evaluator.^8^

Analysis of 162 *de novo* pathogenic CNVs from rare disease patients indicated that Horizon identified 135 as pathogenic variants (83%) and 27 as VUS, achieving an overall AUC of 0.87. This analysis confirmed Horizon’s advantage over AnnotSV and ClassifyCNV, which had AUCs of 0.84 and 0.80, respectively (Figure 1G). The cohort predominantly included deletions (158/214) over duplications (56/214), a known bias on reporting more deletion CNVs as pathogenic. In this cohort of 162 *de novo* pathogenic variants, Horizon accurately identified 135 as pathogenic and 27 as VUS, not misclassifying any as benign. Meanwhile, AnnotSV and ClassifyCNV reclassified 2 and 4 variants as benign, respectively.

Deletions and duplications of this cohort showed equal performance (AUC = 0.87) (Figure 1H), indicating balanced model capability. Size-based analysis revealed the strongest accuracy in CNVs sized 3-5 Mb (AUC = 0.96), followed by those less than 1 Mb (AUC = 0.80) and 1-3 Mb (AUC = 0.71). The least accuracy was observed in CNVs larger than 5 Mb (AUC = 0.50) (Figure 1I).

Across all pathogenic variants analyzed, Horizon’s ROC analysis underscored its robust performance with an AUC of 0.94, distinctly outperforming AnnotSV and ClassifyCNV, which achieved AUCs of 0.85 and 0.81, respectively (Figure 2). This comprehensive evaluation illustrates Horizon’s enhanced reliability in classifying CNV pathogenicity across various cohorts and CNV sizes.

**Figure 2.**
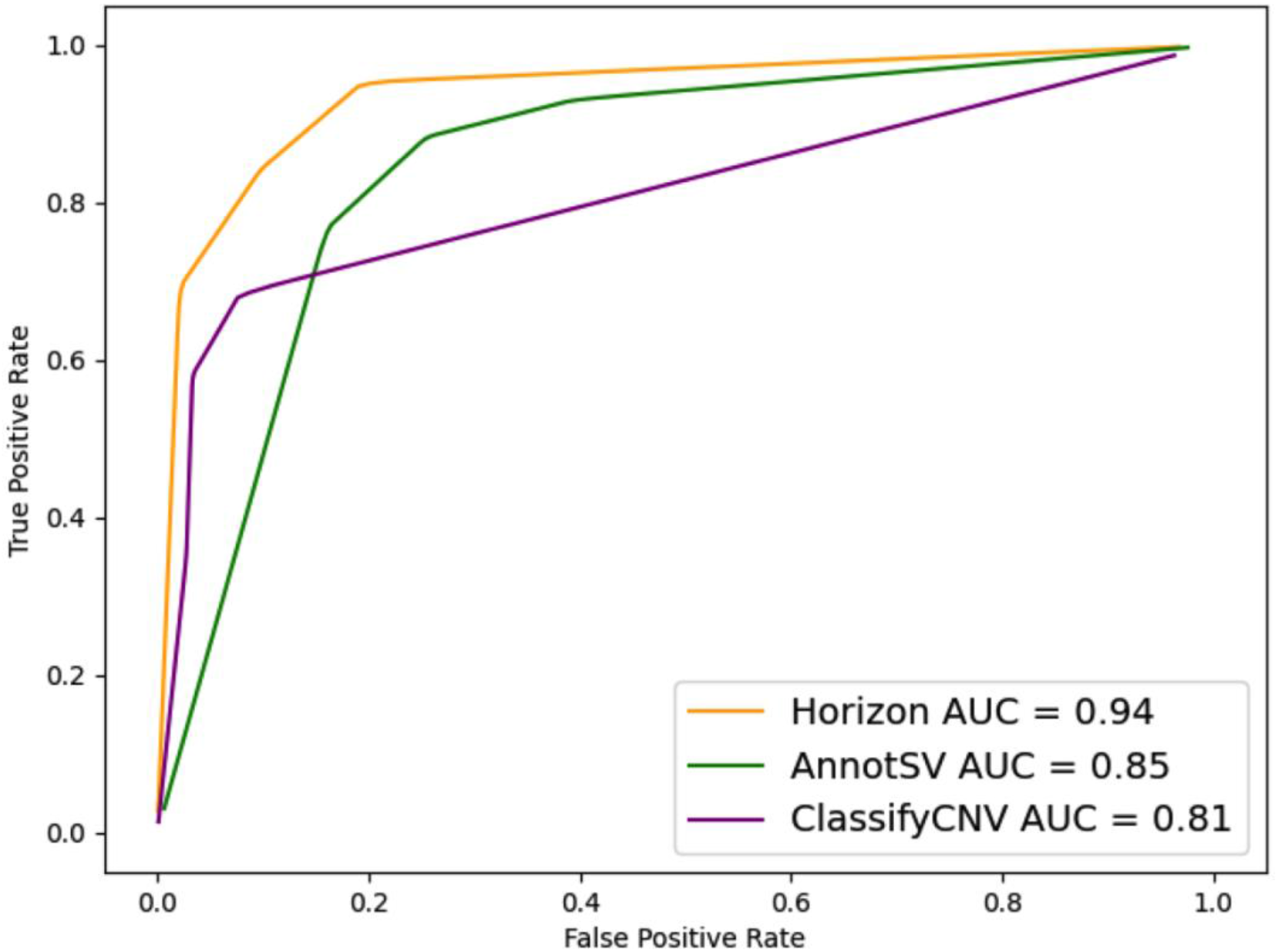
Comparative performance of different tools across all cohorts. A comprehensive summary of the performance comparison between the three tools (Horizon, AnnotSV, and ClassifyCNV) across all cohorts combined. The ROC curve for each tool is color coded to yellow, green, and purple for Horizon, AnnotSV, and ClassifyCNV, respectively.

## Discussion

The rapid development of genome sequencing has enabled increasing insights into genetic variants and their association with various diseases. However, the interpretation of these vast number of genomic data remains a significant bottleneck in the genomic landscape. In this study, we scrutinized the performance of the Horizon model across three datasets comprising CNVs from a discovery cohort and published literature. The Horizon model, based on ACMG guidelines, amalgamates various CNV attributes to ascertain pathogenicity. Utilizing 696 reported pathogenic variants from these cohorts, Horizon’s annotations were benchmarked against clinician interpretations.

Our analysis focused on quantifying the sensitivity and specificity of Horizon’s pathogenicity classification in different CNV datasets. The discovery cohort was crucial in this respect, as the data had less bias and incidental findings were excluded. In this cohort, the Horizon model exhibited an AUC of 0.97, underscoring its accuracy in pathogenicity predictions. Intriguingly, Horizon also demonstrated a marginally superior performance in interpreting duplication CNVs compared to deletions. This finding is notable considering the inherent complexity in predicting the outcomes of duplication CNVs, which are traditionally more challenging to interpret than deletions due to their less predictable effects.^18,19^ The model’s adeptness at discerning smaller CNV sizes, which are typically more difficult to evaluate, further emphasizes its robustness. Horizon’s consistency was evident by its highest accuracy for 3-5 Mb CNVs across all cohorts. However, its performance varied in other length groups depending on the dataset. Interestingly, all pathogenic variants that Horizon failed to recall as pathogenic were subsequently labeled as VUS, not benign. Prediction accuracy was particularly high for the discovery cohort and *de novo* cohort. The literature cohort consists of old microarray and BAC array CNV breakpoints that are widely inaccurate with breakpoints often comprising many critical genes that lead to false pathogenic predictions.^12^ Even in this suboptimal cohort, Horizon’s variant interpretation capabilities outperformed ClassifyCNV and AnnotSV. This superiority is likely attributable to the comprehensiveness of the proprietary databases integral to Horizon’s predictive model. Additionally, Horizon did not reclassify any of the established pathogenic variants as ‘benign,’ in contrast to both ClassifyCNV and AnnotSV.

Horizon stands out due to its utilization of extensive proprietary databases, integrating features like CNV size, type, gene impact, control frequency, and crucially, zygosity. This aspect is particularly relevant as it enables the differentiation of variants based on their potential impact in heterozygous versus homozygous states, a critical factor in the identification of Mendelian disorders such as cystic fibrosis.^20^ The integration of these features, aligned with ACMG guidelines, facilitates the rapid and automated interpretation of variants, addressing a critical need in the genomic field. As such, Horizon enhances the accuracy of genomic interpretations and accelerates the diagnostic process, paving the way for more efficient clinical decision-making and personalized medicine strategies.

## Supporting information

Supplementary Doc

## Data availability

The authors confirm that all data relevant to this study are included within the main text and are further detailed in the Supplementary Figures and Tables accompanying this article.

## Funding Statement

Mohammed Bin Rashid University of Medicine and Health Sciences (MBRU) - College of Medicine, Grant/Award Number: IG-2022; Bangabandhu Science and Technology Fellowship Trust (BSTFT), Grant/Award Number: BDGV-03; AlMahmeed Collaborative Research Awards, Grant/Award Numbers: ALM 1801, ALM 20-0074; Internal grant awards from Genetics and Genomic Medicine Centre, NeuroGen Healthcare, Grant/Award Number: Y2402; Aljalila foundation research grant, Grant/Award Number: AJF2023-207

## Author Contributions

Conceptualization: MU, NN, ME, SS; Data Curation: MU, NN, ME, SS, HA; Formal Analysis: MU, NN, ME, SS, AI, HA, NH, TH; Funding Acquisition: MU; Methodology: MU, NN, ME, SS, AI; Supervision: MU, NN, SH, MW, PR, WK, BB; Writing-original draft: MU, NN, ME, SS; Writing-review and editing: MU, NN, SH, MW, PR, WK, BB

## Conflict of Interest

The authors declare no conflicts of interest.

## Notes

### Competing Interest Statement

The authors have declared no competing interest.

