## Supplementary Doc for "Horizon: CNV interpretation through rapid automated ACMG-aligned pathogenicity analysis"

**Supplementary Tables:**

**Table S1: Cohort details, phenotypic descriptions, and pathogenicity annotations by Horizon model.**

| **Cohort** | **Total Number of CNVs in Cohort** | **Number of Pathogenic CNV’s in Cohort** | **Total Number of Pathogenic CNV’s called by Horizon** | **Phenotype** | **PMID** |
| --- | --- | --- | --- | --- | --- |
| Discovery Cohort | 1,766 | 43 | 77 | NDD | 36959829 |
| Literature Cohort | 708 | 491 | 385 | Developmental delay, Autism, Inborn errors of immunity, Nuchal translucency, oligohydramnios etc. | 37873196, 36553144, 28295210, 28613040, 29693535, 26777411, 31447483, 37895217, 33528536, 33230148, 25106414, 36907537,  30564460, 35486341, 34531397, 31475041, 25333781, 28846756, 37807935, 36180924, 33800913, 29449963, 25937001, 24689080,  37243546, 28771251, 28554332, 32573669, 31564432, 28567303, 31780822 |
| *de novo* Cohort | 214 | 162 | 140 | Congenital heart disease, Cerebral palsy, Intellectual Disability, Eplilepsy, Congenital anamolies etc. | 37873196, 25333781, 31475041, 29449963, 36180924, 33528536, 28613040, 28567303, 31447483, 30847515, 31564432, 28554332,  35506549, 28771251, 30712880, 25205790, 28846756, 31231543, 26777411 |

**Table S2: Horizon predictions on Pathogenic CNV’s across all cohorts.**

| **Horizon Predictions** | **Pathogenic CNV’s** | | | **Total** |
| --- | --- | --- | --- | --- |
|  | **Discovery Cohort** | **Literature Cohort** | ***de novo* Cohort** |  |
| Pathogenic | 41 | 343 | 135 | 519 |
| VOUS | 2 | 148 | 27 | 177 |
| Benign | - | - | - | - |
| Total | 43 | 491 | 162 | 696 |
